# The Digital Mouse: why computational modelling of mouse models of disease can improve translation

**DOI:** 10.1101/2020.05.04.075812

**Authors:** Natal van Riel, Ralph Müller, Enrico Dall’Ara

**Affiliations:** Eindhoven University of Technology, Department of Biomedical Engineering, Eindhoven, the Netherlands; Amsterdam UMC-AMC Campus, Department of Experimental Vascular Medicine, Amsterdam, the Netherlands; Eidgenossische Technische Hochschule Zurich, Institute for Biomechanics, Zurich, Switzerland; University of Sheffield, Department of Oncology and Metabolism, Sheffield, United Kingdom

**Keywords:** regulatory decision-making, computational modelling, *in silico* pre-clinical trial, mouse models, aging, digital twin, geroprotectors, frailty

## Abstract

Computational models can be used to study the mechanistic phenomena of disease. Current mechanistic computer simulation models mainly focus on (patho)physiology in humans. However, often data and experimental findings from preclinical studies are used as input to develop such models. Biological processes underlying age-related chronic diseases are studied in animal models. The translation of these observations to clinical applications is not trivial. As part of a group of international scientists working in the COST Action network MouseAGE, we argue that in order to boost the translation of pre-clinical research we need to develop accurate *in silico* counterparts of the *in vivo* animal models. The Digital Mouse is proposed as framework to support the development of evidence-based medicine, for example to develop geroprotectors, which are drugs that target fundamental mechanisms of ageing.

**Highlights:**

- Computational modelling of human (patho)physiology is advancing rapidly, often using and extrapolating experimental findings from preclinical disease models.
- The lack of *in silico* models to support *in vivo* modelling in mice is a missing link in current approaches to study complex, chronic diseases.
- The development of mechanistic computational models to simulate disease in mice can boost the discovery of novel therapeutic interventions.
- The ‘Digital Mouse’ is proposed as a framework to implement this ambition. The development of a Digital Mouse Frailty Index (DM:FI) to study aging and age-related diseases is provided as an example.

## 1. Introduction

Regulatory bodies require new medical devices and pharmacological interventions to be tested in animals before going to clinical trials. Irrespective of development of advanced *in vitro* systems, such as organoids and organ-on-a-chip, animal models remain part of the regulatory procedure in the foreseeable future. In particular mouse models play a role in the development and implementation of evidence-based interventions. MouseAGE (COST Action BM1402) is a network of academic and industry scientists, clinicians and regulators that aims to reach consensus on ways to test pre-clinical interventions in ageing mice, to improve both relevance and reproducibility of experimental findings, in particular focusing on the multiple facets of frailty [1], [2]. In mice more data on biophysical and molecular processes are available or can be collected than for humans. Machine learning and other computational methods become increasingly important to analyse complex biomedical data. Nevertheless, translation of findings in pre-clinical studies to predict clinical outcome largely relies on the interpretation by researchers. This way of knowledge transfer by human reasoning appears inadequate because of its subjective, non-quantitative and often non-systematic nature. Mathematical models to simulate disease development and treatment response are complementary to data-driven and statistical models [3]. Mechanistic models are based on a mathematical description of a system using mechanical, chemical, physical and biological knowledge about a phenomenon or process. Hence, by definition, these models make use of prior knowledge available in the domain. Simulation models are well-established in engineering sciences and applied in the design and implementation of complex, man-made systems. Computer simulation becomes increasingly important in biomedical research and supports regulatory processes for new medical devices and drugs [4], [5]. However, there is remarkably little attention for a systematic approach to incorporate differences in *in vivo* physiology, disease development and treatment response between humans and animals in the computer simulation models. We argue that *in vivo* modelling using mice can be much more effective when it is supported by specific *in silico* modelling of the manifestation of the disease in the animal and its response to treatment. However, research is required to systematically incorporate differences between the ultimate target, the human patient, and the main source of data and mechanistic understanding, i.e. animals and other pre-clinical models. To fill this gap, we propose the ‘Digital Mouse’ (DM), which is a framework composed of multiple, different mechanistic computational models of mouse (patho)physiology. Using frailty as an example, we outline the opportunities to advance *in silico*, mechanistic modelling to pre-clinical studies, calling for scientists from different fields to work together and in close collaboration with regulatory bodies.

## 2. Virtual Patients in need of a Digital Mouse

Computational modelling is a rapidly developing area in biomedical sciences, and is becoming mature for human physiology (e.g. the VPH, [6]) with applications in understanding disease (Systems Medicine) and development of drugs (Systems Pharmacology) [7]. Mechanistic computer simulation models are applied for hypothesis generation and testing, to support analysis of complex data, to discover biological mechanisms, and modelling contributes to the development of medical therapies. *In silico* we can describe and study different components of disease without confounding factors of cooccurring pathologies (and treatment) that exist in animal models and humans. Such ‘*in silico medicine*’ is already feasible when underlying mechanisms of a disease are (reasonably well) known, and relevant outcomes (biomarkers) are available [8], [9]. For example, mechanistic models of glucose homeostasis are accepted by the FDA as part of the pre-clinical testing for the development of an artificial pancreas [10], [11]. However, similar computer models currently do not exist for complex traits such as age-related chronic diseases (e.g. cardio-metabolic diseases, osteoporosis, frailty).

There exists a rich history of mathematical and computational models to learn about human physiology and disease. Often, these models are based on observations in human studies in combination with data and information from other organisms, such as small animals. (One might call these ‘Frankenstein’ models, alike the creature that was created in the 1818 novel by Mary Shelley.) Though at first sight it seems efficient to direct the *in silico* modelling immediately towards its application for human disease, this approach might actually complicate the translation of pre-clinical findings to clinical research and application [12]. Physiological differences between men and mice can be large and *in vivo* experimental data collected in mice may better *not* be directly used as input for *in silico* models for human applications. Though the *in silico* Frankenstein model built from different data sources may provide insight in generic aspects of biology and disease, it is neither adequately representing the animal, nor is it accurate for humans. Instead, experimental data from mice should better be analysed and integrated with dedicated, mouse-specific *in silico* models. A ‘digital mouse’ next to a ‘virtual patient’ model to facilitate transfer and translation of findings from one species to the other (Figure 1). Hence, to advance clinical trials to study human disease and develop therapy we need *in silico* models of the animal as well.

**Figure 1.**
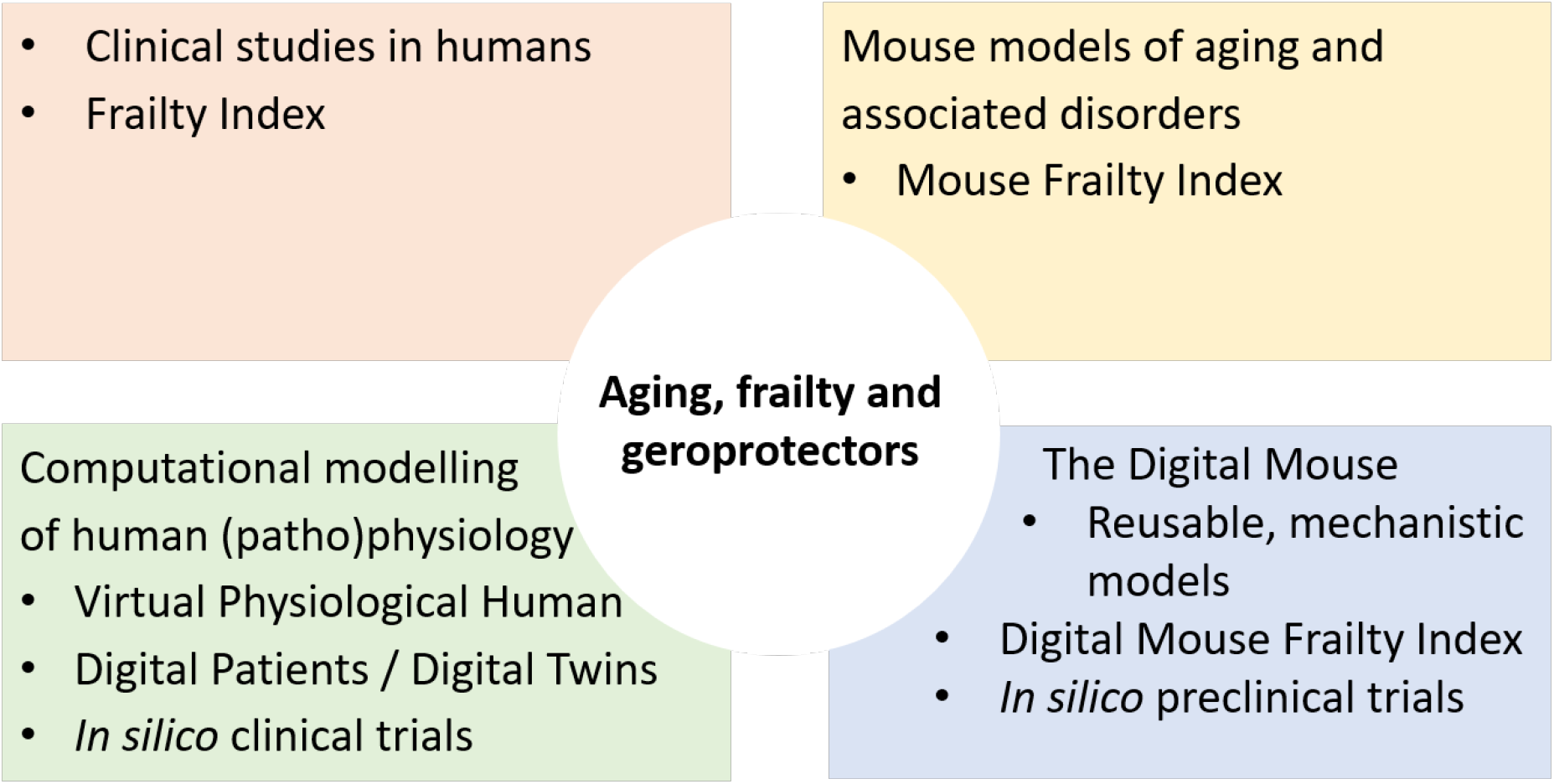
To study complex, chronic age-associated diseases and develop novel therapies the Digital Mouse complements and connects developments in three other domains: human studies, in vivo modelling in mice, and computational modelling of human (patho)physiology.

The envisioned DM is a framework that collects modules and submodels, based on mechanistic descriptions of biology, biophysics and physiology that can be simulated to provide quantitative predictions. It is designed to predict the effect of interventions and to estimate quantities otherwise not measurable noninvasively and longitudinally in the animal. Examples of existing mechanistic computational models specifically for mouse models of disease include metabolism [13] [14] and excitation-contraction coupling in muscle [15]. For bone biomechanics and mechano-biology [16], [17] detailed models can be constructed by combining data from *in vivo* high-resolution imaging [18] with single-cell “omics” technologies [19]. Mechanistic models and simulation techniques to predict disease onset and progression in mice have been developed [20], and were applied to predict the effect of pharmacological interventions in pre-clinical studies with a longitudinal design [21].

The DM is applicable in, what we call, ‘*in silico* pre-clinical trials’ to simulate the effect of treatments, taking into account variability in disease development and treatment response in the *in vivo* model. The term *in silico* pre-clinical trials indicates: *the use of mechanistic computational models in pre-clinical research to simulate treatment responses of in vivo and in vitro experimental systems in the development or regulatory evaluation of medical interventions and biomedical products.* Next to *in silico* trials that use computer simulations to mimic humans [4], *in silico* pre-clinical trials provide a complementary approach to address the imperfection of predictions issued from laboratory and animals studies when applied to humans. Where found effective, *in silico* pre-clinical trials will impact the process, procedures, ethical concerns and costs for the development of new biomedical therapies and products. *In silico* approaches targeted to preclinical studies with animals, in the long term, will contribute to the reduction, refinement and replacement (3R) of animal experiments. Effective deployment of both clinical and pre-clinical *in silico* trials can reduce and partially replace clinical trials as they are being performed today.

## 3. Application in aging and associated diseases: frailty

Geroprotectors are a novel class of drugs that target fundamental mechanisms of ageing and offer an interesting new avenue for therapeutic research [22]. Since it is a new field it could be argued that innovative, *in silico* approaches are more easily adopted compared to disease areas with a long history and established ‘modus operandi’ to develop and evaluate drugs and other interventions. The establishment of aging as a ‘drugable target’ will require basic research involving *in vivo* models (especially mouse models), to identify underlying mechanisms and define endpoints and biomarkers that are translatable to humans [23]. In the development of geroprotectors and other new drugs it is critical to identify off-target effects and other side-effects before first in human trials [24], [25] and develop an understanding of how pharmacokinetic parameters translate from animals to man [26]. Frailty has been conceptualised as a physiological syndrome of decreased reserve and resilience, resulting in progressive functional decline, increased vulnerability to many stressors, and an increase in negative health outcomes and dependence [27]. Frailty is a suitable starting point to develop the Digital Mouse concept in the context of aging and age-associated diseases. Recently, a frailty index (FI) for mice has been presented [28], which is similar to the frailty index for humans according to Rockwood c.s. [29]. The FI considers frailty to be related more to the number rather than to the nature of the health problems, and is defined as the ratio of the number of health deficits accumulated by the individual on the total number of potential deficits evaluated (approximately 30 parameters). This method has been demonstrated to be robust, i.e. not sensitive to the choice of particular items, and to allow the estimation of survival probability without reference to chronological age. For these reasons it has been adopted, with several variations, as a proxy measure of aging and mortality. We can imagine the development of an *in silico* version of the frailty index: the Digital Mouse Frailty Index (DM:FI), which includes attributes that we obtain from simulation of different computational models. Similar to the pre-clinical and clinical FI, the underlying computational models can reflect very different aspects (molecular mechanisms, pathways, tissues and organ systems), of what we consider relevant to describe frailty in a quantitative and predictive manner (Figure 2). Existing computational models and new ones should be simulated in response to multiple stressors and both quantitative and qualitative outcomes should be analysed and transferred into frailty related deficits that can be scored. Existing computational models could be adapted and refined to fit the purpose as submodel of the DM:FI. Computational models of missing parts are to be developed.

**Figure 2.**
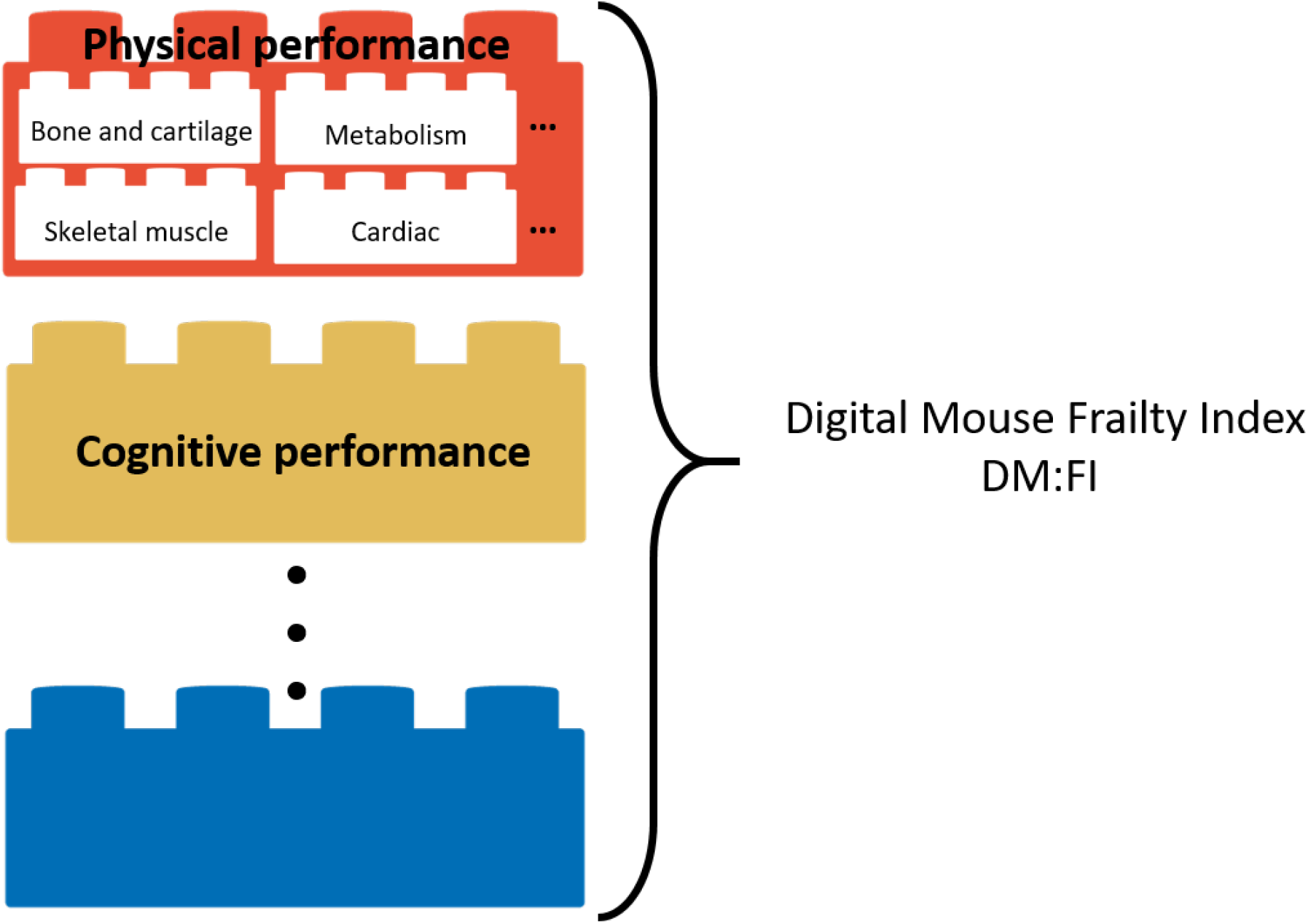
The Digital Mouse Frailty Index (DM:FI) is composed of modules and each module contains several submodels. Modules and submodels can be used and reused in different configurations.

## 4. Outlook

The DM to study ‘virtual’ pathophysiology of chronic age-associated diseases and its application in *in silico* pre-clinical trials to enhance translation offer a unique opportunity for collaborative research at the interface of the physical and life sciences. Endeavours such as outlined here come with many challenges, including technologies needed to build multi-scale and modular models. Another challenge is the development of credible models despite the presence of uncertainty in model components and in the experimental data used for model calibration. Sources of uncertainty include so-called structural uncertainty in the model and uncertainty (variance) in experimental data [30]. Structural uncertainty resides in simplifications that are inherent to the process of model building and assumptions that are made in case molecular mechanisms are unknown or disputed in the field. It is important that models are not only evaluated to describe a certain dataset, but also carry physiological realism [31]. Whereas some model parameters can be directly measured, many will need to be inferred using parameter identification approaches. When model parameters are estimated by calibrating the model to experimental data, uncertainty in the data (noise, errors) will propagate into the parameter estimates

[32], which subsequently will limit the accuracy of model predictions [33], [34]. The complexity of the computational models, the limitations in the amount of experimental data that can be collected per individual animal, and the inherent uncertainties and variance among individuals in the measurements, require a careful strategy to build and test the models, such as the VVQU paradigm (Verification, Validation and Uncertainty Quantification) used by engineers and adopted by the FDA [35]. This is to guarantee that the models are applicable in the context for which they are developed and can make reliable predictions. Development of the *in silico* models requires experimental data of high quality, in particular data collected in longitudinal study designs. It is important that the models describe not only the average disease phenotype in a population, but also account for the heterogeneity to explain variability in treatment responses for individuals. One approach to achieve this is to sample the uncertainty in the model parameters and generate parameter sets for which the model simulations capture the variability in the experimentally observed parameters. Next, the ‘virtual population’ is stratified and divided in subgroups to reflect the distribution of population-level treatment response data [36], [37].

The DM can make use of progression in the field driven by *in silico* modelling of human (patho)physiology. Developments in the field of artificial intelligence (AI) (i.e. machine learning, deep learning) offer opportunities to couple mechanistic and data-driven models. The ‘black box’ methods offered by AI can be used to generate input data for the mechanistic models, or to explain the ‘residuals’, that is information contained in the data that cannot be captured by the mechanistic model. AI, data science and mathematical modelling are also combined in ‘digital twins’, which are computational simulation models of complex systems [38]. In the healthcare domain a digital twin can be a computer model of an individual, that simulates reality (the other twin) and its outputs provide actionable knowledge and facilitate decision making, such as identification of the individually best therapy. Potential applications could be in surgical simulations and virtual reality in computer assisted surgery [39]. The DM provides a framework to develop digital twinning for mice (Figure 3), in which the mouse and the corresponding virtual model are connected by different types of data, and the model automatically updates as the biological counterpart changes. Integration of AI approaches to improve the predictive ability of mechanistic models is a highly relevant topic for future research.

**Figure 3.**
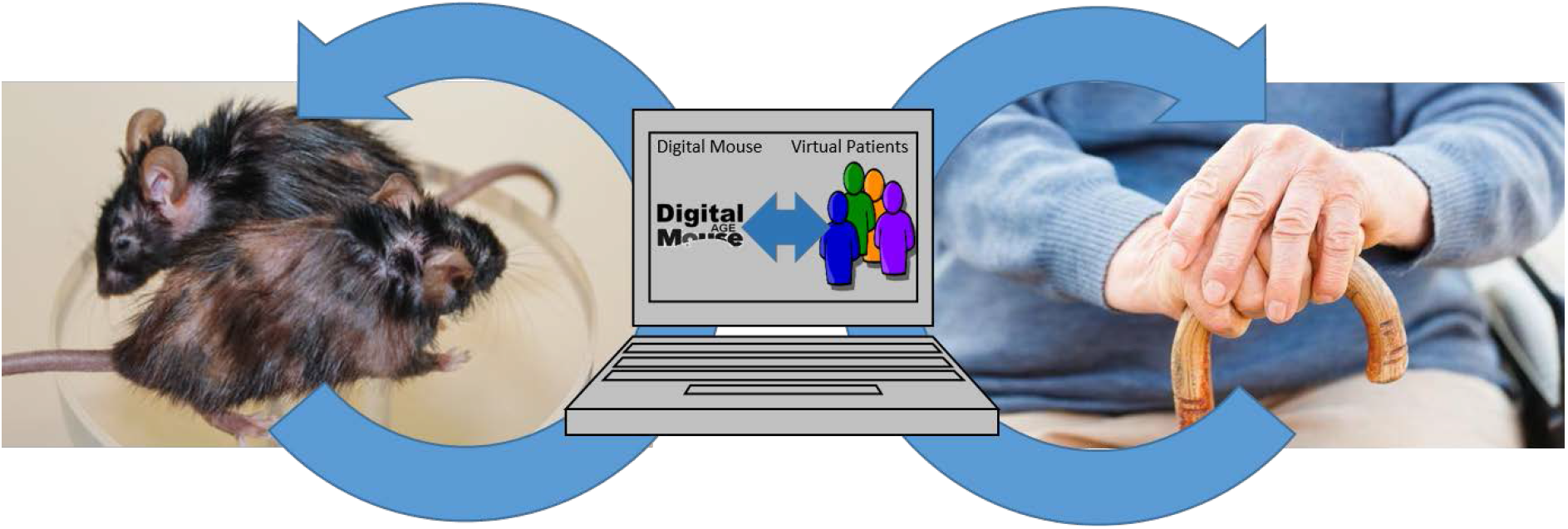
Digital Twinning for mice and humans. The combination of in silico models for the same disease and treatment in both animal models (left) and in humans (right) is expected to improve transfer of knowledge and translatability of pre-clinical data.

We encourage the scientific community to further develop the DM concept, via inclusive and multidisciplinary collaborative networks. This paper could serve as template for grant applications in the European Union, but also world-wide. The development of digital twinning for the life sciences and health and medical care is gaining momentum^1^. The DM could also serve as challenge for scientific team competitions, for example in the form of a ‘modelathon’^2^.

In conclusion, motivated by an increasing role for mechanism-based mathematical models and computer simulations in the approval process for new medical devices and drugs by regulatory bodies, we identify opportunities to systematically develop mechanistic computational models of mouse (patho)physiology. The combination of *in silico* models of response to treatment in both animal models and in humans could drastically improve our accuracy of predicting clinical outcomes from pre-clinical data. The lack of *in silico* models to support *in vivo* modelling in mice is a missing link in current approaches to study aging and age-related diseases. The DM is proposed as concept and framework to improve the translation of preclinical research to successful intervention in human patients. The development of computational models to simulate frailty in mice will contribute to the unveiling of new aspects of the biology of ageing and can boost the discovery of novel therapeutic interventions that prevent frailty and increase resilience to age-related disorders.

## Acknowledgements

This article is based on work from COST Action MouseAGE (BM1402) and was supported by COST (European Cooperation in Science and Technology).

## Footnotes

1 For example, FET Proactive call FETPROACT-EIC-07-2020 Digital twins for the life-sciences, https://ec.europa.eu/info/funding-tenders/opportunities/portal/screen/opportunities/topic-details/fetproact-eic-07-2020, website visited 04/05/2020.

2 A hackathon aimed at modelling, as recently organized by one of the co-authors, http://multisim-insigneo.org/modelathon-2020-optimisation-of-interventions-for-osteoarthritic-patients-with-multi-scale-modelling/, website visited 04/05/2020.

